# Modelling disease transmission from touchscreen user interfaces

**DOI:** 10.1101/2020.10.28.358820

**Authors:** Andrew Di Battista, Christos Nicolaides, Orestis Georgiou

## Abstract

The extensive use of touchscreens for all manner of human-computer interactions has made them plausible instruments of touch-mediated disease transmission. To that end, we employ stochastic simulations to model human-fomite interaction with a distinct focus on touchscreen interfaces. The timings and frequency of interactions from within a closed population of infectious and susceptible individuals was modelled using a basic queuing network. A *pseudo* reproductive number (*R*) was used to compare outcomes under various parameter conditions. We also expanded the simulation to a specific real-world scenario; namely airport self check-in and baggage drop. Results revealed that the required rate of cleaning/disinfecting of screens to effectively mitigate *R* can be inordinately high. This suggests that revised or alternative methods should be considered.

## Introduction

The ubiquitousness of shared Touchscreen User Interfaces (TUIs) has become apparent in recent years; whether it be a fast-food menu or an airport terminal self check-in machine. However, their reputation for hygiene has come under scrutiny, predominantly from sensationalised media articles [1–3]. The fact that touchscreens carry pathogens is not however in question here; what is yet to be established is if they can transmit enough pathogens to a user so as to cause infection (and if so, which disease?), either due to an isolated TUI interaction event, or when thought of as a series of interactions by multiple users, effectively forming an interaction network between them.

Modelling of fomite-mediated disease transmission has already been undertaken in [4, 5], where the authors describe an Environmental Infection Transmission System (EITS) using a system of ODEs incorporating the dominant parameters: pathogen infectivity, survival/persistence on surfaces, and finger-to-surface (surface-to-finger) transfer rates. Other important parameters include the frequency in which people interact/touch the fomite and how often it is disinfected and cleaned [6].

In [7], the authors review the specific role of biometric fingerprint scanners in the transmission of SARS-CoV2. They reiterate the importance of the parameters incorporated in the EITS in addition to recommending enhanced universal hygiene methods; e.g., hand washing, glove wearing, regular surface cleaning and, importantly, the use of non-contact technologies as an overall alternative.

Estimating reasonable values for each disease model parameter relies on often limited experimental and clinical data. For example, the survival rate of pathogens on surfaces has been a source of some contention [8, 9]; quantifying viral particles and bacteria is fundamentally not an exact science [10, 11]. This is further complicated by the reporting of pathogen quantities using incompatible measures, e.g., PFU, CFU, TCID_50_, viral copies from PCR, etc. Moreover, a model needs to consider human behaviour; self-inoculation (transferring of pathogens onto mucosal membranes e.g. mouth, nose, eyes) occurs when individuals touch their faces with contaminated hands. It must be assumed that personal hygiene practices such as handwashing are not strictly nor universally adhered to.

With all of these considerations, rather than target a specific disease model (e.g., Influenza, SARS-CoV2, etc.), in this paper we focus on establishing/reasserting the fundamental mathematical parameters that govern fomite-mediated disease transmission and examine how each can affect outcomes. We use stochastic Monte-Carlo simulations as they offer more flexibility and ease in incorporating the large number of parameters versus traditional ODE analysis [5, 12–15].

With specific regard to TUIs, we also examine the impact of using touchless technologies and alternatives such as computer hand tracking using cameras, proximity sensors, RADAR, mid-air haptics [16] and other ‘touch-free’ interface solutions and compare their effectiveness to the current leading alternative i.e. more frequent cleaning/disinfection.

## Materials and methods

### Basic scenario and assumptions

For transmission via fomite to occur an Infectious donor (*I*_1_) must first interact with the fomite and deposit some amount of pathogens onto its surface. Pathogens must then survive long enough for a sufficient dose to be subsequently picked-up by the hands/fingers of a Susceptible (*S*) host. This newly Exposed (*E*) individual must then transfer these pathogens onto the mucosal membrane regions of their face (e.g., eyes, mouth, nose), i.e., self-inoculate if they are to become Infected (*I*_2_) [17].

When considering shared TUIs such as those found in public spaces, we can make some further general assumptions:

- Individuals will use the TUI in sequence (i.e. they behave as if in a queue).
- Given typical touchscreen menu design, users are obliged to touch the same regions of the screen e.g. confirmation buttons, on-screen keypads etc. Therefore, regardless of the application or screen size, users are essentially *sharing the same surface area*.
- We can assume that all touch events are carried out with finger tips (possibly just the index finger of the dominant hand).
- For transmission to occur we assume individuals are *not* washing their hands before/after using the interface.
- An infectious person can be defined as someone who has relatively high initial levels of pathogens on their hands at any given time e.g. from coughing/sneezing into one’s hands or a having recently used the toilet without hand-washing afterwards. *They remain infectious throughout the simulation*.
- Once a susceptible person becomes exposed (*S* → *E*), we assume self-inoculation (face touching) occurs within 20 minutes after TUI use. Beyond a certain time it is unreasonable to attempt to model the level of pathogens on a finger; exposed individuals will have interacted with countless other fomites and surfaces (e.g.wiping hands on clothing, opening doors, using their mobile phones, etc.). There is also the issue of pathogen survival on skin that can range from a few minutes to several hours [18–20]. The 20 minute assumption mitigates the effects of these uncertainties. Therefore, the exposed state is transitory: An exposed individual will either self-inoculate (*E* → *I*_2_) or revert back to a susceptible state (*E* → *S*) after TUI use.
- For the same reasons as above, we do not model the re-deposition of pathogens from one TUI to another (when users interact with more than one TUI). Infected individuals do not contribute to pathogen deposition; they carry on, for all intents and purposes, as susceptible individuals except we do not count or consider their additional self-inoculation events.
- In all simulations, we consider a single time period (i.e., a day) using a 1-minute time-step. We do not consider incubation periods, or recovery rates. This is because many pathogens may lead to infection (but do not subsequently render the exposed person *infectious*). (See Outcome measures).

The three main actors in this scenario are therefore the *pathogens*, the network of *touchscreens* and the *network of people*, each having its own controlling parameters. The remainder of this section describes the implementation of this computer model and present pseudo code where applicable to provide the reader with the best clarity. A summary table listing all relevant parameters and description can be found in Appendix: Table A.1.

Monte-Carlo simulations generally make use of a variety of sampling distributions in order to model random events. Throughout this paper we use of the *rate parameter λ* to describe a *Poisson* process and the symbol *p* to describe its discrete time counterpart, the *Bernoulli* process. When considering random variables sampled over a particular range, [*a, b*], we make use of the *truncated normal distribution*, denoted *f* (*x*; *μ, σ, a, b*), where *x* is a random variable with mode *μ* and variance *σ*^2^. Other random variables are sampled from uniform distributions, *U* [*a, b*].

### Queuing network model

The movement of people is simulated using a system of first-in-first-out (FIFO) queues (Fig. 1). We begin with an initial population pool of *N* = (*S* + *I*_1_) people (we can assume there is an existing disease prevalence *I*_1_*/N*). These individuals leave the pool at a rate *λ*_0_, and arrive at one (or any) of *L locations*. Each location has an *arrival* and *departure* queue. Arrivals are people who have yet to interact with the TUI, departures are those who have already interacted and are ready to (potentially) move on to another location (into that location’s *arrival* queue). *Locations* can be interpreted as places where there is a cluster of identical TUIs e.g. a kiosk of ATM machines. An establishment may have several locations within it, each with a different number and type of TUIs serving different customer functions.

**Fig 1.**
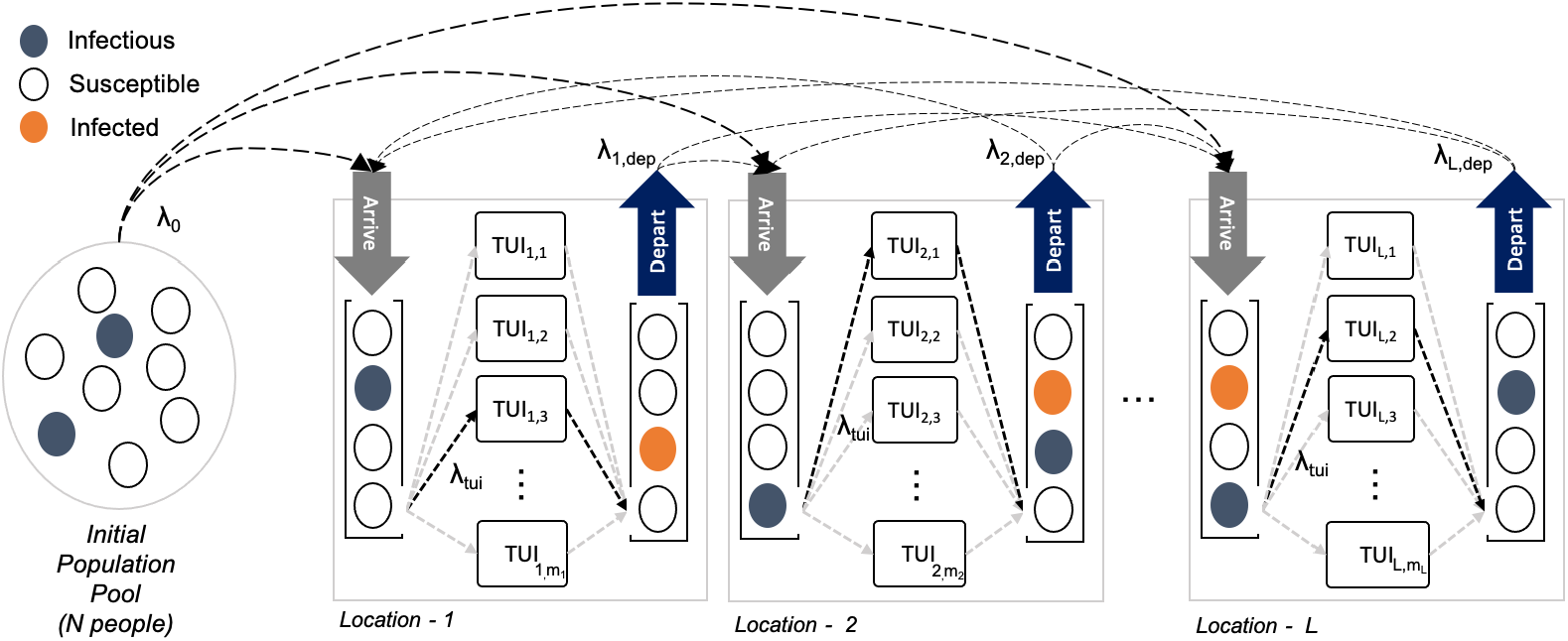
Queuing network. Infectious and susceptible people from an initial population pool enter a new location at a rate *λ*_0_. The location model consist of an *arrival* and *departure* FIFO queue; people in the arrival queues wait to interact with one of a number of TUIs, they then move on to the *departure* queue and (potentially) move on to another location at rate *λ*_dep_. Markov-Chains of conditional probabilities along with a parameter to set the total number of ‘jumps’ from one location to another is used to govern the flow of people. The rate at which people move through the queues is dictated by the number of TUIs at each location and the rate parameter, *λ*_tui_.

The arrival queues are depleted at a rate *m*_*j*_λ_*tui*_, (*j* = 1, 2, … , *L*), where *m_j_* is the number of TUIs at the *j^th^* location and *λ*_tui_ is the rate of TUI use i.e. 1/*λ*_tui_ is the average time interval between TUI use. In our simulations *λ*_tui_ is kept constant across all locations so that a TUI is used on average once every 2 minutes (in general it is a parameter associated with each individual TUI design). After a TUI interaction, people may stay at that location for some time before moving on, e.g., eating at a fast food restaurant after ordering a meal. The rate at which people *depart* the location is governed by *λ*_dep_. Subsequent movement of people between locations is controlled via a Markov Chain of conditional probabilities and a *number-of-jumps* parameter that sets how many locations a person can visit before being removed from the active simulation. This framework allows for modelling anything from very basic to increasingly elaborate networks of people movements.

### Touchscreen model

#### Transfer efficiency asymmetry

An important assumption in this model is that susceptible (*S*), exposed (*E*) and newly infected (*I*_2_) individuals can only *pick-up* pathogens from a TUI surface. Deposition of pathogens onto TUIs is carried out by infectious donor (*I*_1_) individuals exclusively. This assumption is also linked to the concept of *transfer efficiency asymmetry*.

Let us define the deposit rate (*α*) as the proportion of pathogens on an infectious finger transferred onto the surface of a fomite. Similarly, we define pick-up rate (*β*) as the proportion of pathogens on a fomite that are transferred to the finger of a susceptible person. With regards to fingers and non-porous surfaces (like glass), transfer efficiency has been shown to be asymmetric. From glass-to-finger, pick-up rates, *β*, are on the order of 20 ± 30(*SD*)% [21, 22], while deposit rates, *α*, are considerably lower i.e. 5% [23, 24] (Fig. 2). Some key points worth noting about the experiments conducted to ascertain these values: deposit rates were measured by inoculating a finger with a known concentration of pathogens and measuring the amount left behind on a clean surface. Pick-up rates were examined by touching a contaminated surface with a clean finger and measuring the pathogen level on the finger. This would suggest that we can interpret the transfer efficiency data as a *one-way* or net result i.e. either pathogen are deposited by an infectious person or picked up by a susceptible, exclusively.

**Fig 2.**
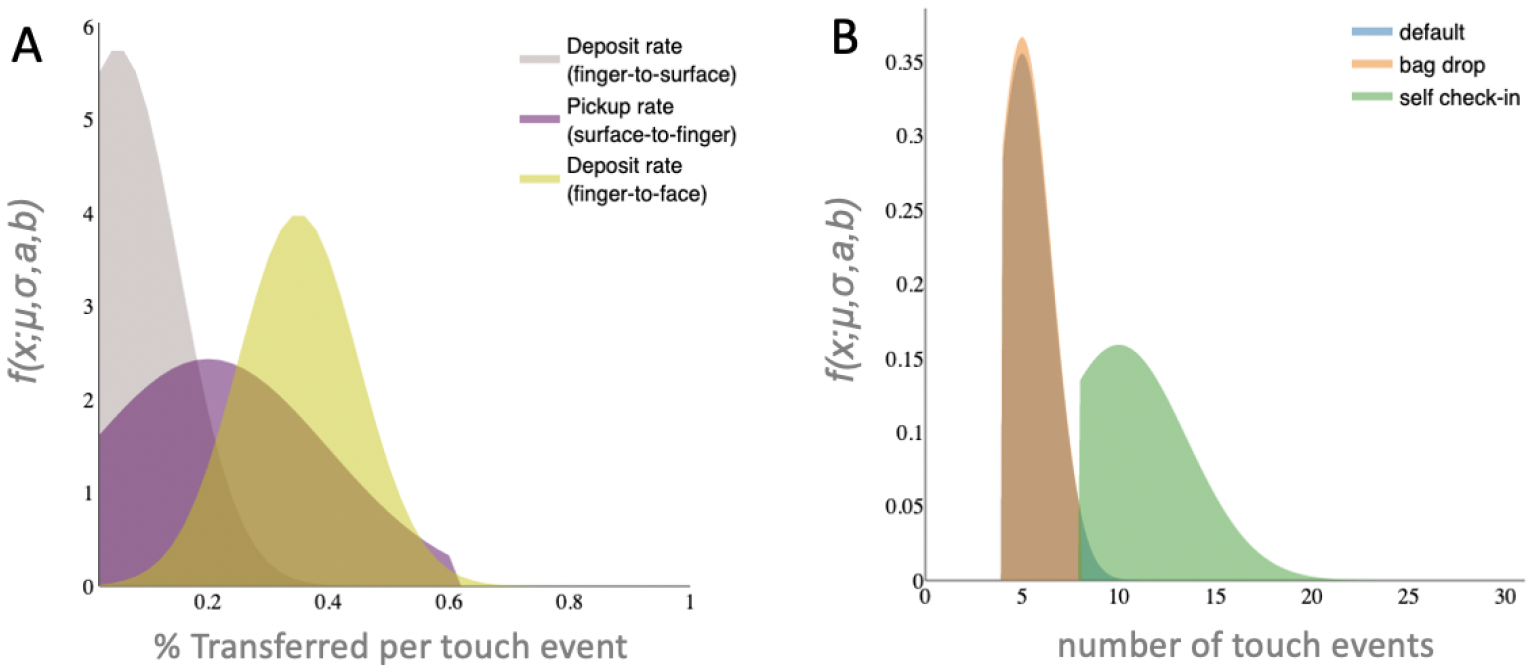
Transfer and touch rates. (A) Pathogen transfer (pick-up (*β*) and deposit (*α*) rates are estimated by sampling from truncated normal distributions; default values estimated from the literature are depicted for surface-to-finger, finger-to-surface and finger-to-face. (B) Using truncated normal distributions to simulate the average number of touch events during TUI interaction; depicted are a ‘default’ model used for simulations along with that of a bag-drop and self check-in machine found at most major airports. To inform our choice of distribution parameters, we made use of online video demonstrations self-check in procedures; advertised by many major airline companies.

We model transfer rates using truncated normal distributions, default values are depicted in (Fig. 2A)). This allows for incorporating the mode, *μ*, and variance, *σ*^2^, (i.e., uncertainty) from experimental findings described in the literature while incorporating bounding limits, (*a, b*), (i.e., 0 to 100%).

#### Touch rates

As with transfer rates, the number of touches, *n*_*t*_, expected to complete a transaction or menu selection can be modelled using a truncated normal distribution. For example, an ATM pin pad requires a minimum of 5 touches (4 pin numbers + OK), i.e., both mode and minimum can be set to 5. However, cancelled transactions, re-attempts, correcting invalid input etc. means that we can expect a small variance in touch numbers. Unless we have compelling reason not to, the maximum number of touches can be safely limited to some reasonably large value, e.g., 30. Fig. 2B depicts the distributions sampled from to model touchscreen events in this simulation; a default *generic* TUI *f* (*n*_*t*_; 5, 1.5, 4, 30), an airport *self check-in machine f* (*n*_*t*_; 10, 3.5, 8, 30) and *bag-drop* kiosk *f* (*n*_*t*_; 5, 1.5, 4, 8).

#### Pathogen removal

Assuming effective cleaning practices, a thorough wipe with an appropriate cleaning agent will remove approximately 98% of pathogens [25]. We simulate cleaning events with a daily frequency *p*_clean_. This removal of pathogens is in conjunction with deactivation / die-off rates of bacteria and viruses which can live on surfaces for several hours to months [26, 27]. We model this using the pathogen half-life on a surface, *t*_1/2_. These parameters are likely play an important role if they are on the same order of magnitude as the intervals between TUI use.

#### Touchscreen dynamical model

Let *D* be the total accumulated level (or dose) of pathogens on a TUI. Alg 1 describes the flow of pathogens at each simulation time step. Note, TUI interaction (and all the calculations to go with it) happens in a single simulation time-step.

**Algorithm 1.**
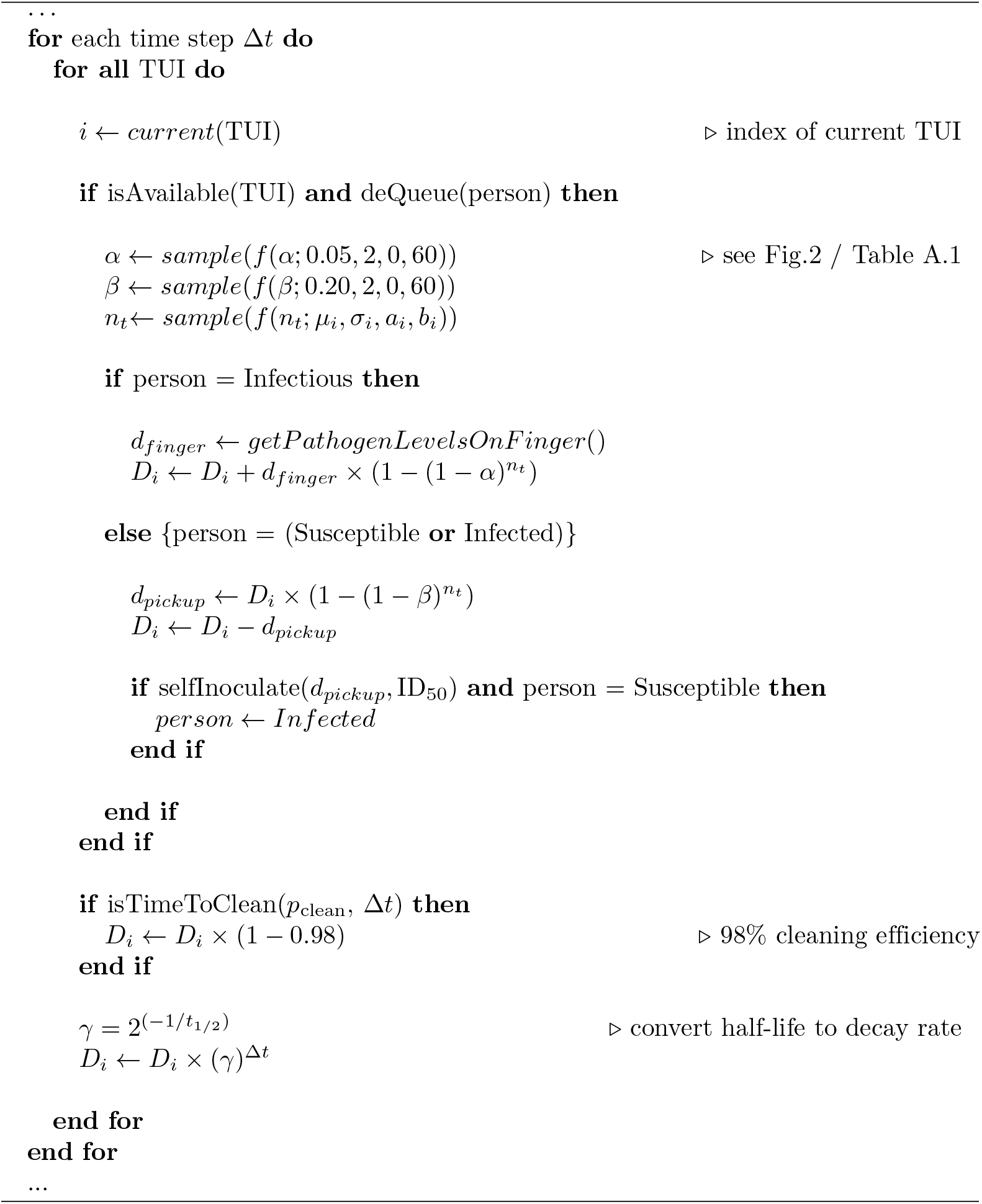
Calculate flow of pathogens in time-step Δ*t*

In the case of an infectious user, *d_finger_* is the number of pathogens *on* their finger. For susceptible users, *d_pickup_* is the number of pathogens transferred *to* their finger. For each interaction we sample a random number of touch events *n*_*t*_ and select a *β* or *α* rate from its respective distribution. Note, we only select a transfer rate once per interaction (technically it should vary with every touch). Here, we assume that an individual will be consistent in the pressure applied with their fingers with each touch (and their fingers will obviously not change in surface area), so transfer rates can be taken as constant throughout the interaction. Computational efficiency savings is another motivating factor.

There are two functions presented in Alg 1 that still have yet to be defined *selfInoculate()* and *getPathogenLevelOnFinger()* which will be discussed in the following sections.

### Pathogen shedding

In our model we would like to estimate the pathogenic load on a *single finger* by considering the following scenarios:

1. After toilet use (taking into account hand washing rates and effectiveness); we may consider both respiratory and enteric viruses (and bacteria).
2. Coughing and/or sneezing into one’s hands (assumes a respiratory virus e.g. influenza)

Viral loads in faeces can be as high as 10^7^ − 10^8^ PFU/g [17, 28]; this can include common respiratory viruses like influenza in addition to enteric disease causing pathogens. The level of Norovirus in faeces has been reported as 10^5^ − 10^9^ particles/g based on PCR [29]. Bacterial load found on hands after toilet use ranged from 0.85 ± 0.93(*SD*)×10^5^CFU for washed and dried hands to 3.64±4.49(*SD*) × 10^5^ CFU for unwashed hands [30]. In other experiments, 10^8^ CFU/g has been used to approximate natural bacterial contamination levels [31].

It has been estimated that 30% of individuals do not wash their hands sufficiently [32] and there are additional issues in lavatories with regards to using contaminated soap [33] and doorknobs [34]. Therefore, we can feel justified in our assumptions about the prevalence of *infectious* individuals in a given population.

The average volume of a cough has been reported at between 0.006 - 0.044 ml [4, 6]. Sneeze volumes are estimated as 40 times that of a cough. Coughing and sneezing rates of influenza sufferers are on the order of 12-22/h and 5/h, respectively. The concentration of viral particles in ex-pulsed droplets, based on nasal swabs, ranges on the order of 10^4^ − 10^5^ TCID_50_/*ml*.

Regardless of the units quoted (TCID, PFU etc.) the *number* of units are ostensibly of similar orders of magnitude. As will be discussed (see Dose response), this number will ultimately be normalised relative to the infectivity of the pathogen under consideration.

From [35] we can estimate that a single fingertip represents approximately 1.4% of the hand’s surface (and shares that proportion of pathogens). Based on all of the above, we estimate the total dose shed from the finger of an infectious individual (prior to each interaction with a TUI) by Alg 2.

**Algorithm 2.**
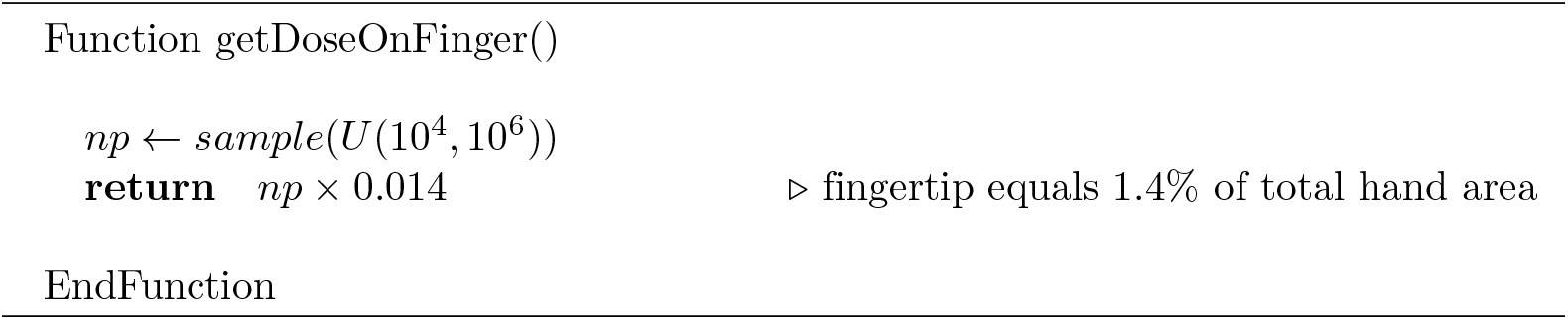
getDoseOnFinger() subroutine

### Self-inoculation

Face touching rates involving direct contact with mucosal membranes (eyes, nose, mouth etc.) have been found to be approximately 15 touches/h [36, 37]. We can simulate the number of face touching events, *k*, in our prescribed 20 minutes period by drawing from a Poisson distribution, Pois(*k*; *λ*), where *λ* = 5 touches every 20 minutes.

The deposit rate of pathogens from finger-to-lip (more generally skin-to skin) has been reported in the region of 35% [24, 36] (see Fig. 2 and Appendix Table A.1).

#### Dose response

The human ID_50_ is the infective dose with 50% probability of infection. Typically, respiratory viruses require a relatively large dose for infection (10^3^ − 10^4^ TCID_50_) [4, 6, 38]. For many types of bacteria and enteric viruses this can be as low as 10 - 100 PFU (or CFU) [39, 40]. If we interpret these values as estimates for ID_50_, provided we stick with the same units, pathogen levels can effectively be *normalised*. It is customary to model a dose response using an exponential cumulative distribution function (CDF) [5]. Self-inoculation is therefore calculated by Alg 3.

**Algorithm 3.**
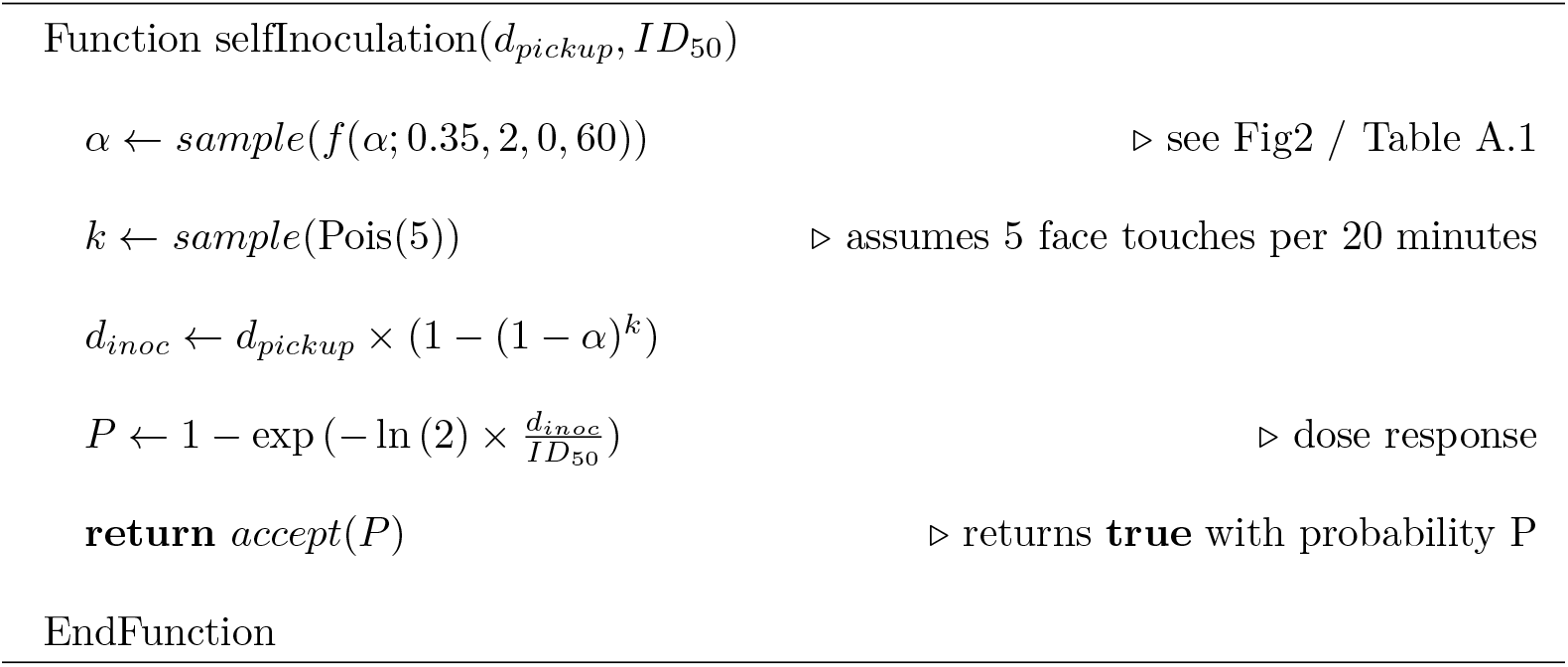
selfInoculation() subroutine

#### Outcome measures

In fomite-mediated transmission a *pseudo reproduction number*, *R* can be defined as the number of susceptible people that the *fomite* infects having been contaminated by an infectious person. Thus, *R* is defined as the ratio of newly infected individuals to the initial number of infectious (Eq 1).

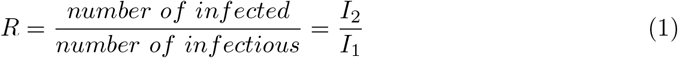

Another metric of interest is the *gap* (in terms of number-of-users) between infectious contamination and subsequent susceptible users becoming infected. It seems intuitive that the next susceptible user will be the most likely to become infected. However, we can measure and store this gap from the simulation results to confirm this assertion. Accordingly, the questions we would like to answer are the following:

- What is the probability of becoming infected after using a TUI?
- On average, how many susceptible individuals could become infected as a direct result of a single infectious user over the course of a day, i.e. *R*?
- Which TUI users are getting infected, i.e. what is the time gap between infectious and infected?
- What is the efficacy of frequent cleaning on reducing the probability of infection?

## Results

### Overview

In this section we present the results from two simulated scenarios; default simulation parameters are listed in Table 1. The touch and transfer rates used are those already discussed and depicted in Fig. 2. Each scenario is simulated over the period of a single day with one minute time-step resolution. Note that care needs to be applied when considering parameters such as population *N*, *λ*_0_, *λ*_tui_, etc. to ensure that the entire population actually makes it through the simulation during the allotted time. Results for each parameter setting are averaged over 10,000 realisations of a single day period.

**Table 1.** Simulation default parameters.

| Parameter | Simulation 1 | Simulation 2 |
| --- | --- | --- |
| Number of locations | 1 | 2 |
| TUIs per location | 1 | (36, 24) |
| $N$ | 100 | 12000 |
| $\lambda_0$ (per minute) | 0.5 | 20 |
| $\lambda_{\text{dep}}$ (per minute) | 0.05 | 1000 |
| $\lambda_{\text{tui}}$ (per minute) | 0.5 | |
| Disease Prevalence | 2% |  |
| ID<sub>50</sub> | 10<sup>2</sup> |  |
| $t_{1/2}$ (hours) | 3 | |
| $p_{\text{clean}}$ (daily) | 0 | |

1. **Simulation 1**: A location with a single TUI; this allows us to examine the model’s sensitivity to parameters (survival rates, infectious dose, cleaning rates, etc.). We also look at the effects of adding extra TUIs at that location.
2. **Simulation 2**: A real-world example involving two TUI locations; Airport terminal *check-in machines* followed by *baggage drop*. We use data for London Heathrow (LHR) Terminal 5 [41] along with the assumption that one in four outgoing passengers makes use of those machines.

Default parameters for Simulations 1 and 2. Simulations are carried out using 10,000 realisations of a 1 day period (with 1 minute time-steps resolution). Simulation 1 models 1 location with a single TUI. Simulation 2 models an airport terminal with 36 self check-in machines and 24 bag-drop machines. The figure of *N* = 12000 is derived from passenger arrival data from LHR T5 (2018) and assuming 1 in 4 passengers actually makes use of the machines [41].

### Simulation 1: a single TUI location

The following figures show the effects on the reproduction number *R* for varying different simulation parameters; *disease prevalence* (Fig. 3A), *pathogen survival*, *t*_1/2_ (Fig. 3B), *infectivity*, ID_50_ (Fig. 3C), the *number of* TUIs (Fig. 3D), *touch rates* (Fig. 3E), *cleaning rate*, *p*_clean_ (Fig. 3F) and the effect of *additional locations* (Fig. 4). For each simulation, we keep all other parameters constant as given in Table 1.

**Fig 3.**
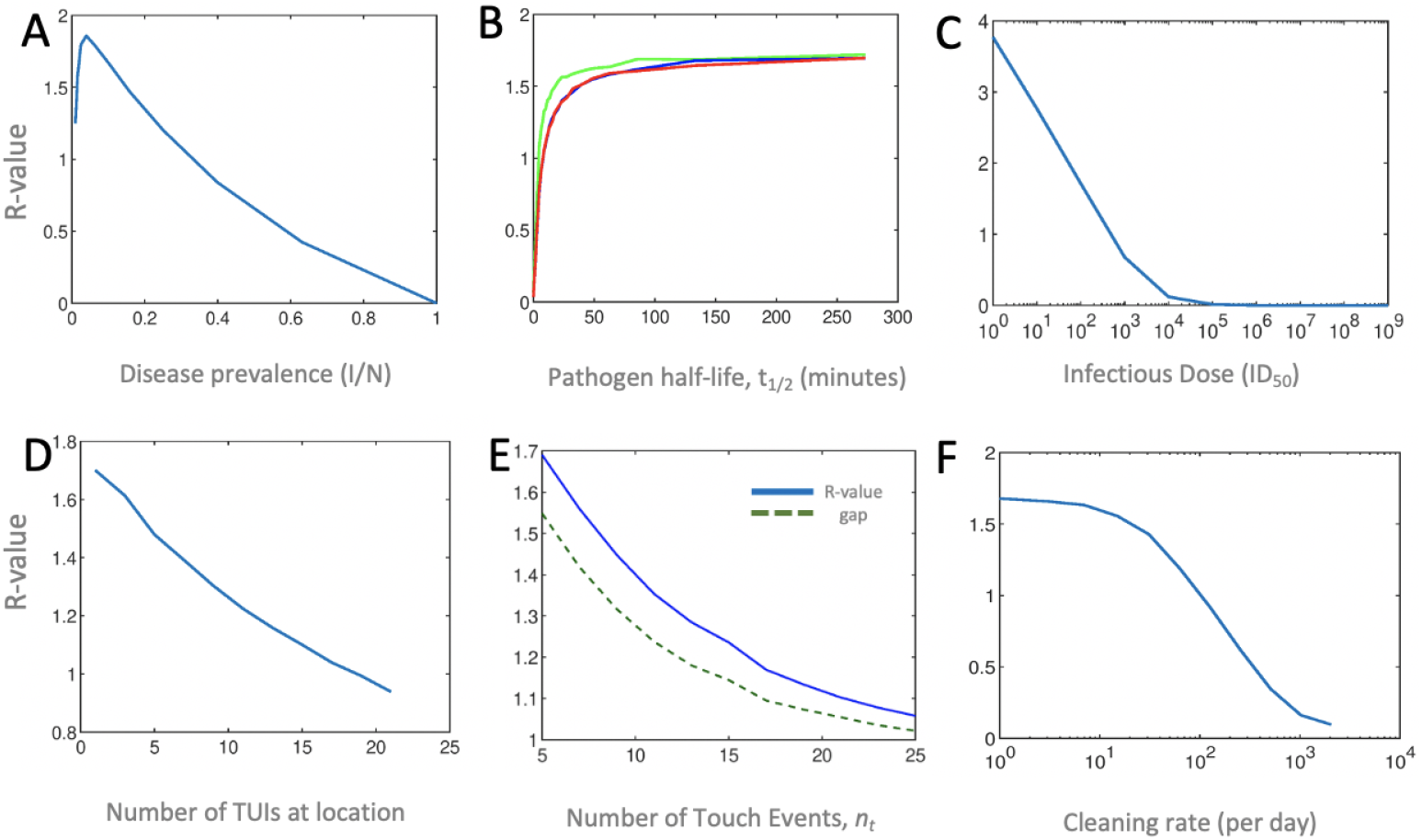
Simulation 1 control parameters. (A) Disease Prevalence: *R*, reaches a peak at approximately 4%, suggesting an optimal cumulative effect of infectious donor prevalence, beyond which it drops steadily along with the proportion of susceptible individuals. (B) Pathogen Survival: A longer half-life results in *R* asymptotically approaching a maximum value of 1.7. Three separate plots were made using *λ*_tui_ values of 0.5, 0.2 and 0.1, i.e., 2 (green), 5(blue) and 10 (red) minutes average interval between TUI use. Longer intervals allows for more time for pathogens to die-off thus slightly lowering the R value. (C) Infectious Dose: The effect of varying parameter ID_50_ on *R* is significant; beyond a certain level of infectivity, fomite-mediated disease transmission becomes non-viable. (D) Number of TUIs: As the number of TUIs available for use at a location increases, the risk of infection drops steadily (approximately linearly). With more TUIs to choose from, the time intervals between their use increases and the effective pick-up and self-inoculation probabilities diminish. (E) Number of touch events: Increasing the average number of touches per TUI interaction lowers the infection rates (blue-solid). The average gap (green-dotted) between infectious and infected users indicates that at increasing touch-rates the *next* susceptible user of a TUI after its contamination (gap=1) almost exclusively becomes infected. In other words, a higher touch rate results in a greater pick-up of pathogens, effectively cleaning the surface for subsequent users (shielding them) while simultaneously increasing the probability of infection for the current user. (F) Cleaning rate: Cleaning rates, *p*_clean_, on the order of several hundred times per day are required to achieve an *R* value less than one. The cleaning intervals are random and are thus not correlated to the rate of TUI users. Therefore low cleaning rates do not effectively prevent the next susceptible user of the TUI from picking-up pathogens.

**Fig 4.**
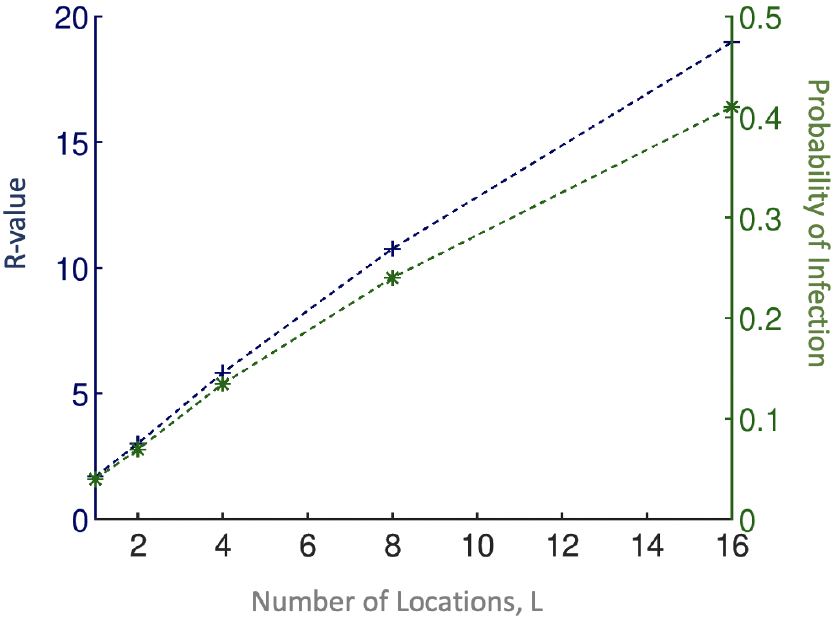
Additional locations. Here we model the effects of additional, identical locations; each simulated person has equal probability of visiting any one location, and will make as many ‘jumps’ as there are locations. Plotted alongside *R* (blue +) is the probability of infection (green *), which is calculated as *I*_2_/*S*. Extra locations means more chances for the infectious donor to spread disease. The plots are not precisely linear; the ratio of *R* to the number of locations *decreases* with *increasing* location numbers. In all simulations, people can only be *infected* once (despite picking-up more infectious doses). Therefore, this reduction in infection rate efficiency is due to herd-immunity.

### Simulation 2: airport terminal with two TUI locations

For this simulation, we focus on the effects of *cleaning rate*, *p*_clean_ (Fig. 5A) and compare its effectiveness to substituting a proportion of TUIs with a ‘touch-free’ alternative (Fig. 5B).

**Fig 5.**
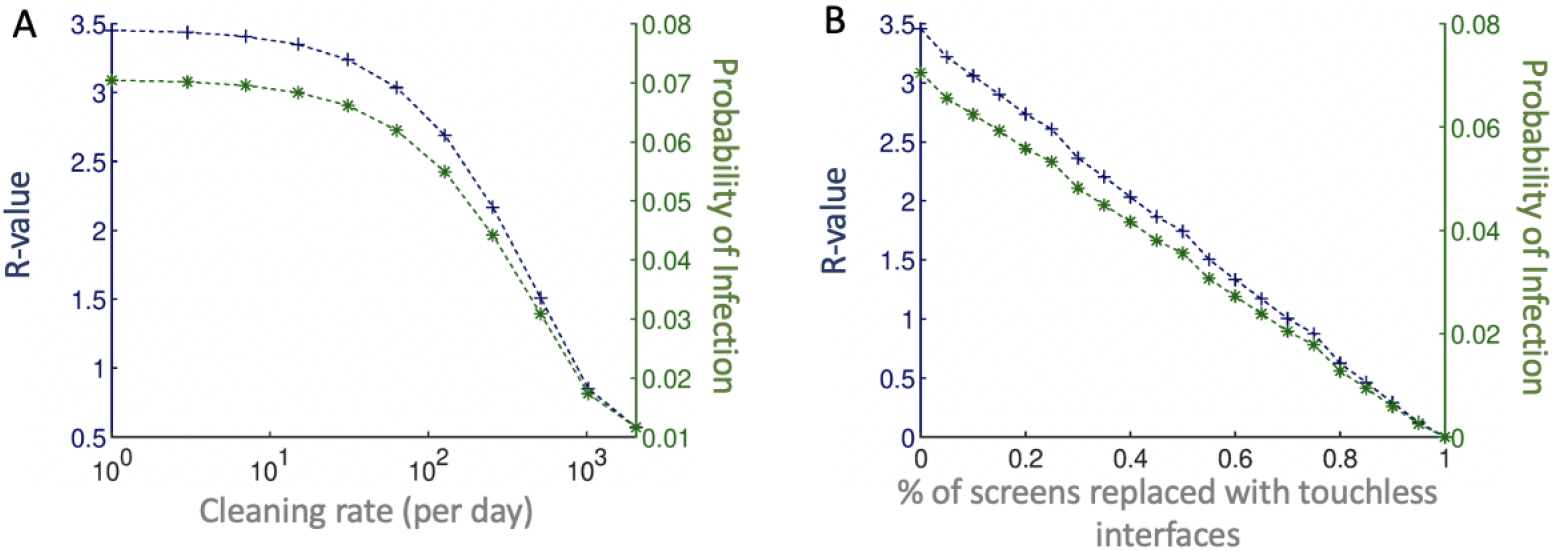
Cleaning rate vs ‘touch-free’ interventions. (A) The effects of cleaning are similar to that in Simulation 1; *R* values are higher overall due to having more than one location. (B) Replacing a proportion of TUIs with a ‘touch-free’ alternative results in a direct linear drop in *R*. For comparison, the same *R* value is achieved by replacement of 50% of TUIs as a cleaning rate of several hundred cleans per day per TUI, of which there are 60 in this simulation.

## Discussion

In many cases the simulation results are intuitive. For example, it is clear that timing plays an important role as the number of TUIs per location (Fig. 3D), pathogen survival (Fig. 3B) and the rate of TUI use, *λ*_tui_, all interact to affect the infection rate. Other unsurprising results are the effects of increasing initial disease prevalence (Fig. 3A) and infectious dose (Fig. 3C).

Increasing the number of locations (Fig. 4), essentially gives infectious individuals multiple chances to contaminate TUIs and infect other users. Were it not for the effects of herd-immunity (and the fact that infections were tallied only in the first instance), *R* values would rise directly proportionally to the number of TUI locations in the simulation. The assumption that an infectious user *always* has some dose of pathogens to deposit at each interaction (Alg. 2), is probably not justified at much higher location numbers as it supposes an inexhaustible shedding of pathogens (or exceptionally unhygienic behaviour). Therefore, Fig. 4 likely overestimates *R* as the number of locations increases.

Rather less intuitive is the effect of increasing the average number of touches, *n*_*t*_, for TUI interaction (Fig. 3E). Throughout all simulations, the average *gap* recorded between infectious and infected user was between 1 and 2; this implies that the susceptible users who *immediately follow* an infectious user are most at risk. Because of the asymmetrical way pick-up and deposit rates are modelled, higher touch rates do the same effective job as cleaning the TUI; the next susceptible user is essentially doomed to infection while simultaneously shielding subsequent users.

In order to use this simulator to model a specific disease, one would have to collate shedding rates and infectious dose information. This requires care when dealing with TCID_50_, PFU, CFU, etc. For example, the default simulation parameters could have been attributed to certain strains of E. Coli or adenovirus. Enteric disease causing pathogens ostensibly have the right combination of (relatively) long half-life on surfaces and low infectious dose to be the major players in fomite-mediated transmission. *Drawing any conclusions about a specific pathogen using this model should be done with caution*.

We did not model the *re-deposition* of pathogens from newly infected individuals onto fomites due to the complexity and consequent unreliability in estimating pathogen levels on an individual’s hand over time. Though unconfirmed, it is reasonable to assume that re-deposition would likely increase the overall infection rates in the scenarios tested.

In this paper we made use of a pseudo *R* value. It should be clear that, while an R value less than 1 is desirable in a pandemic, in this scenario a user-interface designer should be aiming much lower (*R* ≪ 1). One effective way to mitigate infection spread via fomites is with compulsory handwashing. This has also been shown to be effective against pandemics in airport networks [15, 42]. Indeed, the relevance of our paper’s results depends on whether or not a population will maintain these stringent hygienic practices.

An alternative approach that places the responsibility and control with the TUI owners/operators is enhanced sterilization regimens. From Fig. 3F and Fig. 5A, it is apparent that cleaning rates on the order of hundreds of times per day per TUI are required to have a significant effect on *R*. From a cost perspective, this can be prohibitive. In addition to the added cost of cleaning agents, protective equipment (e.g. gloves) and increased staff exposure, the excessive use of industrial and household cleaning agents carries with it a health risk, particularly to those with breathing ailments [43, 44].

UV light is an alternative to chemical agents for disinfecting surfaces, but does not solve the issue of increased cleaning rates nor is it completely absolved from health implications [45–48].

A promising and attractive solution is the emerging technology of self-cleaning antimicrobial surface coatings, many of which are commercially available e.g. for tablets/smartphones [49–51]. With respects to the model presented in this paper, these coatings would have significant impact on the pathogen half-life parameter, *t*_1/2_, (Fig. 3B). However, not all pathogens are significantly affected [52]. It is also unclear if their antimicrobial properties diminish with regular use or require maintenance and/or replacement.

Another alternative to enhanced cleaning are touch-free interfaces that completely remove the need to touch and therefore deposit or pickup pathogens from surfaces (Fig. 5B). Whether or not businesses and venues choose to implement such an alternative will likely depend on the cost of replacement or conversion of existing TUIs. They also need to consider human/consumer behaviour and expectations, particularly in times of pandemics, regarding hygiene. What is for certain however, is that the COVID-19 pandemic has already forced a massive world-wide digital revolution in how we live, work, and do business.

## Acknowledgments

This research project was conducted in collaboration with Ultraleap Ltd. and the authors gratefully acknowledge their advice and support. Orestis Georgiou has received funding from the European Union’s Horizon 2020 research and innovation programme under the Marie Sklodowska-Curie project NEWSENs, grant agreement No. 787180. Christos Nicolaides has received funding from the European Union’s Horizon 2020 research and innovation programme under the Marie Sklodowska-Curie project NISIHealth, grant agreement No. 786247.

## Code availability

The code that supports the findings of this study is available as open-source at https://gitlab.com/fomite-simulator/fomite_sim.

## Appendix

**Table A.1.** Simulation parameters.

| Symbol | Description | Distribution/Range |
| --- | --- | --- |
| **Pathogen Model** |  |  |
| $n_p$ | Number of pathogens shed by an infectious individual | $U(10^4, 10^6)$ |
| $ID_{50}$ | Infectious Dose (50%. Probability) | $10^0 - 10^9$ |
| $t_{1/2}$ | Pathogen half life on surface | 5 minutes – 24 hours |
| **Self Inoculation Model** |  |  |
| $k$ | Face touch events per 20 minutes | $Pois(k; 5)$ |
| $\alpha_{face}$ | Deposit rate (finger to face) | $f(\alpha_{face}; 0.35, 0.1, 0, 1)$ |
| **TUI Model** |  |  |
| $\alpha_{tui}$ | Deposit rate (tui) | $f(\alpha_{tui}; 0.05, 0.1, 0, 0.6)$ |
| $\beta$ | Pick-up rate (tui) | $f(\beta; 0.20, 0.2, 0, 0.6)$ |
| $n_t$ | Number of touches (default) | $f(n_t; 5, 1.5, 4, 30)$ |
| | Number of touches (self-checkin) | $f(n_t; 10, 3.5, 8, 30)$ |
| | Number of touches (bagdrop) | $f(n_t; 5, 1.5, 4, 8)$ |
| $\lambda_{tui}$ | TUI use rate (per minute) | 0.5 |
| $p_{clean}$ | TUI cleaning rate (daily) | 0 - 1440 |
| **Queuing Network Model\*** |  |  |
| $N$ | Population | 100 - 12000 |
| $\lambda_0$ | Initial arrival rate (per minute) | 0.2 - 20 |
| $\lambda_{dep}$ | Departure rate (per minute) | 0.05 - 1000 |

Summary of simulation parameters. \*Not listed are the number of locations, TUIs per location and disease prevalence ( $I_1/N$ ) associated with the queuing network model.

## References

1. Healthline. Want to Avoid Dangerous Bacteria? Don’t Use Public Touch Screens; 2018. Available from: https://www.healthline.com/health-news/want-to-avoid-dangerous-bacteria-dont-use-touch-screens.

2. Health M. Traces Of Poo Have Been Found on Every McDonald’s Touchscreen; 2019. Available from: https://www.menshealth.com/uk/health/a759768/traces-of-poo-found-on-every-mcdonalds-touchscreen-kiosk-tested/.

3. Post TW. No, McDonald’s touch screens are not contaminated with poop; 2018. Available from: https://www.washingtonpost.com/world/2018/11/29/no-mcdonalds-touch-screens-are-not-contaminated-with-poop/.

4. Li S, Eisenberg JNS, Spicknall IH, Koopman JS. Dynamics and Control of Infections Transmitted From Person to Person Through the Environment. American Journal of Epidemiology. 2011;170:257–265. doi:10.1093/aje/kwp116.

5. Zhao J, Eisenberg JE, Spicknall IH, Li S, Koopman JS. Model Analysis of Fomite Mediated Influenza Transmission. PLoS ONE. 2012;7:257–265. doi:10.1371/journal.pone.0051984.

6. Zhang N, Li Y. Transmission of Influenza A in a Student Office Based on Realistic Person-to-Person Contact and Surface Touch Behaviour. Int J Environ Res Public Health. 2018;15. doi:10.3390/ijerph15081699.

7. Okereafor K, Ekong I, Markson IO, Enwere K. Fingerprint Biometric System Hygiene and the Risk of COVID-19 Transmission. JMIR Biomedical Engineering. 2020;5. doi:10.2196/19623.

8. Musa EK, Desai N, Casewell MW. Potential role of inanimate surfaces for the spread of coronaviruses and their inactivation with disinfectant agents. Infection Prevention in Practice. 2020;2. doi:10.1016/j.infpip.2020.100044.

9. Goldman E. Exaggerated risk of transmission of COVID-19 by fomites. Lancet Infect Dis. 2020;doi:10.1016/S1473-3099(20)30561-2.

10. Smithera SJ, Lear-Rooney C, Biggins J, Pettitt J, Jr GGO, Lever MS. Comparison of the plaque assay and 50assay as methods for measuring filovirus infectivity. Journal of Virological Methods. 2013;193:565–571. doi:10.1016/j.jviromet.2013.05.015.

11. l Goldman E. An Overview of Virus Quantification Techniques. Virocyt; 2013. Available from: https://pdfs.semanticscholar.org/a634/aaf93397c2eb7a14af6b1ba1ee5232d1b92d.pdf.

12. Harma G, MacLehose R, Richardson D. Markov chain Monte Carlo: an introduction for epidemiologists. International Journal of Epidemiology. 2013;42:627–634. doi:10.1093/ije/dyt043.

13. Cauchemez S, Carrat F, Viboud C, Valleron AJ. A Bayesian MCMC approach to study transmission of Influenza: application to household longitudinal data. Statistics in Medicine. 2004;23:3469–3487. doi:10.1002/sim.1912.

14. Cauchemez S, Carrat F, Viboud C, Valleron AJ. Bayesian inference of epidemiological parameters from transmission experiments. Nature: Scientific Reports. 2017;7. doi:10.1038/s41598-017-17174-8.

15. Nicolaides C, Avraam D, Cueto-Felgueroso L, González MC, Juanes R. Hand-Hygiene Mitigation Strategies Against Global Disease Spreading through the Air Transportation Network. Risk Analysis. 2020;40(4):723–740. doi:10.1111/risa.13438.

16. Carter T, Seah SA, Long B, Drinkwater B, Subramanian S. UltraHaptics: multi-point mid-air haptic feedback for touch surfaces. In: Proceedings of the 26th annual ACM symposium on User interface software and technology; 2013. p. 505–514.

17. Otter JA, Donskey C, Yezlic S, Douthwaited S, Goldenberg SD, Weber DJ. Transmission of SARS and MERS coronaviruses and influenza virus in healthcare settings: the possible role of dry surface contamination. Journal of Hospital Infection. 2016;92:235–250. doi:10.1016/j.jhin.2015.08.027.

18. Ansari SA, Sattar SA, Springthorpe VS, Wells GA, Tostowaryk W. Rotavirus Survival on Human Hands and Transfer of Infectious Virus to Animate and Nonporous Inanimate Surfaces. Journal of Clinical Microbiology. 1988;26:1513–1518.

19. L’Huillier AG, Tapparel C, Turin L, Boquete-Suter P, Thomas Y, Kaiser L. Survival of rhinoviruses on human fingers. Clinical Microbiology and Infection. 2015;21. doi:10.1016/j.cmi.2014.12.002.

20. Musa EK, Desai N, Casewell MW. The survival of Acinetobacter calcoaceticus inoculated on fingertips and on formica. Journal of Hospital Infection. 1990;15:219–227. doi:10.1016/0195-6701.

21. Lopez GU, Gerba CP, Tamimi AH, Kitajima M, Maxwell SL, Roseb JB. Transfer Efficiency of Bacteria and Viruses from Porous and Nonporous Fomites to Fingers under Different Relative Humidity Conditions. Applied and Environmental Microbiology. 2013;79:5728–5734. doi:10.1128/AEM.01030-13.

22. Julian TR, Leckie JO, Boehm AB. Virus transfer between fingerpads and fomites. Journal of Applied Microbiology. 2010;109:1868–1874. doi:10.1111/j.1365-2672.2010.04814.x.

23. Greene C, Ceron NH, Eisenberg MC, Koopman J, Miller JD, Xi C, et al. Asymmetric transfer efficiencies between fomites and fingers:Impact on model parametrization. American Journal of Infection Control. 2018;46:620–626. doi:10.1016/j.ajic.2017.12.002.

24. Greene C, Eisenberg MC, Vadlamudi G, Foxman B. Fomite-fingerpad transer efficieny(pick-up and deposit) of Acinetobacter baumannii - With and without a latex glove. American Journal of Infection Control. 2015;36. doi:10.1016/j.ajic.2015.05.008.

25. Lopez GU, Kitajima M, Havas A, Gerba CP, Reynolds KA. Evaluation of a Disinfectant Wipe Intervention on Fomite-to-Finger Microbial Transfer. Applied and Environmental Microbiology. 2014;80:3113–3118. doi:10.1128/AEM.04235-13.

26. Boone SA, Gerba CP. Significance of Fomites in the Spread of Respiratory and Enteric Viral Disease. Applied and Environmental Microbiology. 2007;73:1687–1696. doi:10.1128/AEM.02051-06.

27. Kramer A, Schwebke I, Kampf G. How long do nosocomial pathogens persist on inanimate surfaces? A systematic review. BMC Infectious Diseases. 2006;130. doi:10.1186/1471-2334-6-130.

28. Vasickova P, Verani M, Pavlik I, Carducci A. Issues Concerning Survival of Viruses on Surfaces. Food and Environmental Virology. 2010;doi:10.1007/s12560-010-9025-6.

29. Teunis P, Vennema H, Sukhrie FHA, Bogerman J. Shedding of norovirus in symptomatic and asymptomatic infections. Epidemidiology and Infection. 2014;143:1–8. doi:10.1017/S095026881400274X.

30. Mutters R, Warnes SL. The method used to dry washed hands affects the number and type of transient and residential bacteria remaining on the skin. Journal of Hospital Infection. 2019;101:408–413. doi:10.1016/j.jhin.2018.12.005.

31. Gerhardts A, Hammer TR, Balluff C, Mucha H, Hoefer D. A model of the transmission of micro-organisms in a public setting and its correlation to pathogen infection risks. Journal of Applied Microbiology. 2012;112:614–621. doi:10.1111/j.1365-2672.2012.05234.x.

32. Lee MS, Hong SJ, Kim YT. Handwashing with soap and national handwashing projects in Korea: focus on the National Handwashing Survey, 2006-2014. Epidemiology and Health. 2015;37. doi:10.4178/epih/e2015039.

33. Zapka CA, Campbell EJ, Maxwell SL, Gerba CP, Dolan MJ, Arbogast JW, et al. Bacterial Hand Contamination and Transfer after Use of Contaminated Bulk-Soap-Refillable Dispensers. Applied and Environmental Microbiology. 2011;77:2898–2904. doi:10.1128/AEM.02632-10.

34. Alwis WRD, Pakirisamy P, San LW, Xiaofen EC. A Study on Hand Contamination and Hand Washing Practices among Medical Students. ISRN Public Health. 2012;doi:10.5402/2012/251483.

35. Goker P, Bozkir MG. Determination of hand and palm surface areas as a percentage of body surface area in Turkish young adults. Trauma Emerg Care. 2017;2. doi:10.15761/TEC.1000135.

36. Nicas M, Best D. A Study Quantifying the Hand-to-Face Contact Rate and Its Potential Application to Predicting Respiratory Tract Infection. Journal of Occupational and Environmental Hygiene. 2008; p. 347–352. doi:10.1080/15459620802003896.

37. Kwok YLA, Gralton J, McLaws ML. Face touching: A frequent habit that has implications for hand hygiene. American Journal of Infection Control. 2015; p. 112–114. doi:10.1016/j.ajic.2014.10.015.

38. Belshe R, Voris LV. Cold-recombinant influenza A/California/10/78 (H1N1) virus vaccine (CR-37) in seronegative children: infectivity and efficacy against investigational challenge. Journal of Infectious Disease. 1984;149:735–740. doi:10.1093/infdis/149.5.735.

39. Schmid-Hempel P, Frank SA. Pathogenesis, Virulence, and Infective Dose. PLoS Pathogens. 2007;3. doi:10.1371/journal.ppat.0030147.

40. Todd E, Greig J, Bartleson C, Michaels B. Outbreaks where food workers have been implicated in the spread of foodborne disease. Part 4. Infective doses and pathogen carriage. J Food Prot. 2008;71:2339–2373. doi:10.4315/0362-028x-71.11.2339.

41. Ltd HA. About Heathrow: Facts and Figures; 2018. Available from: https://www.heathrow.com/company/about-heathrow/facts-and-figures.

42. Nicolaides C, Cueto-Felgueroso L, González MC, Juanes R. A metric of influential spreading during contagion dynamics through the air transportation network. PloS one. 2012;7(7):e40961.

43. Zock J. World at work: Cleaners. BMJ: Occupational and Environmental Medicine. 2004;62. doi:10.1136/oem.2004.015032.

44. Chang A, Schnall AH. Cleaning and Disinfectant Chemical Exposures and Temporal Associations with COVID-19. Weekly. 2020;69:496–498.

45. Muzslay M, Yui S, Ali S, Wilson APR. Ultraviolet-C decontamination of hand-held tablet devices in the healthcare environment using the Codonics D6000™ disinfection system. Journal of Hospital Infection. 2018;100(3):e60–e63. doi:https://doi.org/10.1016/j.jhin.2018.04.002.

46. Ontario HQ. Portable Ultraviolet Light Surface-Disinfecting Devices for Prevention of Hospital-Acquired Infections: A Health Technology Assessment. Ontario Health Technolgy Assessment Series. 2018;18(1):1–73.

47. Memarzadeh F, Olmsted RN, Bartley JM. Applications of ultraviolet germicidal irradiation disinfection in health care facilities: Effective adjunct, but not stand-alone technology. American Journal of Infection Control. 2010;38(5, Supplement):S13–S24. doi:https://doi.org/10.1016/j.ajic.2010.04.208.

48. Surdu S, Fitzgerald EF, Bloom MS, Boscoe FP, Carpenter DO, Haase RF, et al. Occupational Exposure to Ultraviolet Radiation and Risk of Non-Melanoma Skin Cancer in a Multinational European Study. Plos ONE. 2013;.

49. McCoy CP, O’Neil EJ, Cowley JF, Carson L, Baroid ATD, Gdowski GT, et al. Photodynamic Antimicrobial Polymers for Infection Control. Plos ONE. 2014;doi:10.1371/journal.pone.0108500.

50. Wei X, Yang Z, Tay SL, Gao W. Photocatalytic TiO2 nanoparticles enhanced polymer antimicrobial coating. Applied Surface Science. 2014;290:274–279. doi:https://doi.org/10.1016/j.apsusc.2013.11.067.

51. Page K, Correia A, Wilson M, Allan E, Parkin IP. Light-activated antibacterial screen protectors for mobile telephones and tablet computers. Journal of Photochemistry and Photobiology A: Chemistry. 2015;296:19–24. doi:https://doi.org/10.1016/j.jphotochem.2014.08.011.

52. Raza I, Raza A, Razaa SA, Sadar AB, Qureshi AU, Talib U, et al. Surface Microbiology of Smartphone Screen Protectors Among Healthcare Professionals. Cureus. 2017;9. doi:10.7759/cureus.1989.

